# Aging-associated alterations in the mammary gland revealed by single-cell RNA sequencing

**DOI:** 10.1101/773408

**Authors:** Carman Man-Chung Li, Hana Shapiro, Christina Tsiobikas, Laura Selfors, Huidong Chen, G. Kenneth Gray, Yaara Oren, Luca Pinello, Aviv Regev, Joan S. Brugge

## Abstract

Aging of the mammary gland is closely associated with increased susceptibility to diseases such as cancer, but there have been limited systematic studies of aging-induced alterations within this organ. We performed high-throughput single-cell RNA-sequencing (scRNA-seq) profiling of mammary tissues from young and old nulliparous mice, including both epithelial and stromal cell types. Our analysis identified altered proportions and distinct gene expression patterns in numerous cell populations as a consequence of the aging process, independent of parity and lactation. In addition, we detected a subset of luminal cells that express both hormone-sensing and alveolar markers and decrease in relative abundance with age. These data provide a high-resolution landscape of aging mammary tissues, with potential implications for normal tissue functions and cancer predisposition.

## Introduction

The mammary gland is a dynamic organ that undergoes constant changes in its cellular composition and gene expression over a lifetime. While most transcriptomic studies on the mammary gland to-date have focused on embryonic development, puberty, and pregnancy^1-3^, relatively little is known about their alterations associated with aging^4-7^, which is an important aspect of developmental biology closely related to diseases such as cancer. Furthermore, in human studies the intrinsic effects of aging are often intertwined with extrinsic effects from environmental factors and lifestyle that are difficult to control. A systematic analysis of the cellular and transcriptomic alterations in aged mammary glands in a well-controlled setting could provide important insights into their normal function and disease susceptibility.

The mammary gland is composed of both epithelial and stromal cells^8^. The epithelium contains two distinct layers, an inner layer of luminal cells and an outer layer of myoepithelial/basal cells. The luminal cells can be further subdivided into two subtypes: hormone-sensing (HS) luminal cells capable of responding to hormonal cues such as estrogen, progesterone, and prolactin, as well as secretory alveolar (AV) luminal cells capable of milk production. In contrast, the myoepithelial cells are distinguished by their contractility for dispersing milk and by their ability to deposit extracellular matrix proteins to the basement membrane. These mammary epithelial cells are surrounded by a diversity of stromal cells, including fibroblasts, vascular/lymphatic endothelial cells, immune cells, and adipocytes. Together, the epithelial and stromal compartments of the mammary gland enable the proper function this complex organ. While recent single-cell transcriptomic profiling efforts have provided valuable insights into multiple mammary epithelial cell types^1-3,9^, a similar analysis of the stromal compartment has not been reported to date.

In this study, we utilized scRNA-seq to quantitatively capture the aging-associated alterations in cell type composition and transcriptomic landscape of the mammary epithelium and stroma, using mouse as a model organism. The use of murine mammary glands allows for precise control of biological factors such as genetic background, parity, and hormone cycle, all of which can significantly affect the state of the mammary gland and are difficult to control for using human samples. Our high-resolution transcriptomic profiling by scRNA-seq uncovered a variety of age-related alterations that would be masked in traditional bulk analyses.

## Results

### Diverse epithelial and stromal cell types in the mammary gland

To study epithelial and stromal cells in young and aged mammary glands, we performed scRNA-seq analysis using the 10X Chromium platform on a cohort of young mice (3-4 months old, n = 3) and aged mice (13-14 months old, n = 4) (Fig. 1a). All animals were of the same genetic background, nulliparous, and analyzed at diestrus to minimize any confounding effects caused by differences in these biological factors. We chose diestrus to facilitate the detection of progenitor cells, as epithelial cell proliferation peaks during maximal progesterone levels at the luteal diestrus phase^9-11^. Fresh enzymatically dissociated single cells were analyzed by scRNA-seq without prior cell type-specific isolation or depletion, thereby allowing us to capture both epithelial and stromal cells. Filtering at the cell and gene levels resulted in a final set of 19,161 cells, with approximately 1,600-4,000 cells from each sample, covering a total of 17,509 genes. Single cells were clustered based on their gene expression profiles using Seurat and visualized using t-distributed stochastic neighbor embedding (t-SNE)^12,13^(Fig. 1b). Scrublet^14^ estimation of cell doublets revealed that they only accounted for 1.25% of our data and did not drive cell clustering, indicating their minimal impact on downstream analyses (Supplementary figure 1a). We then identified the specific epithelial and stromal cell types using canonical gene markers (Fig. 1b-f).

**Fig. 1.**
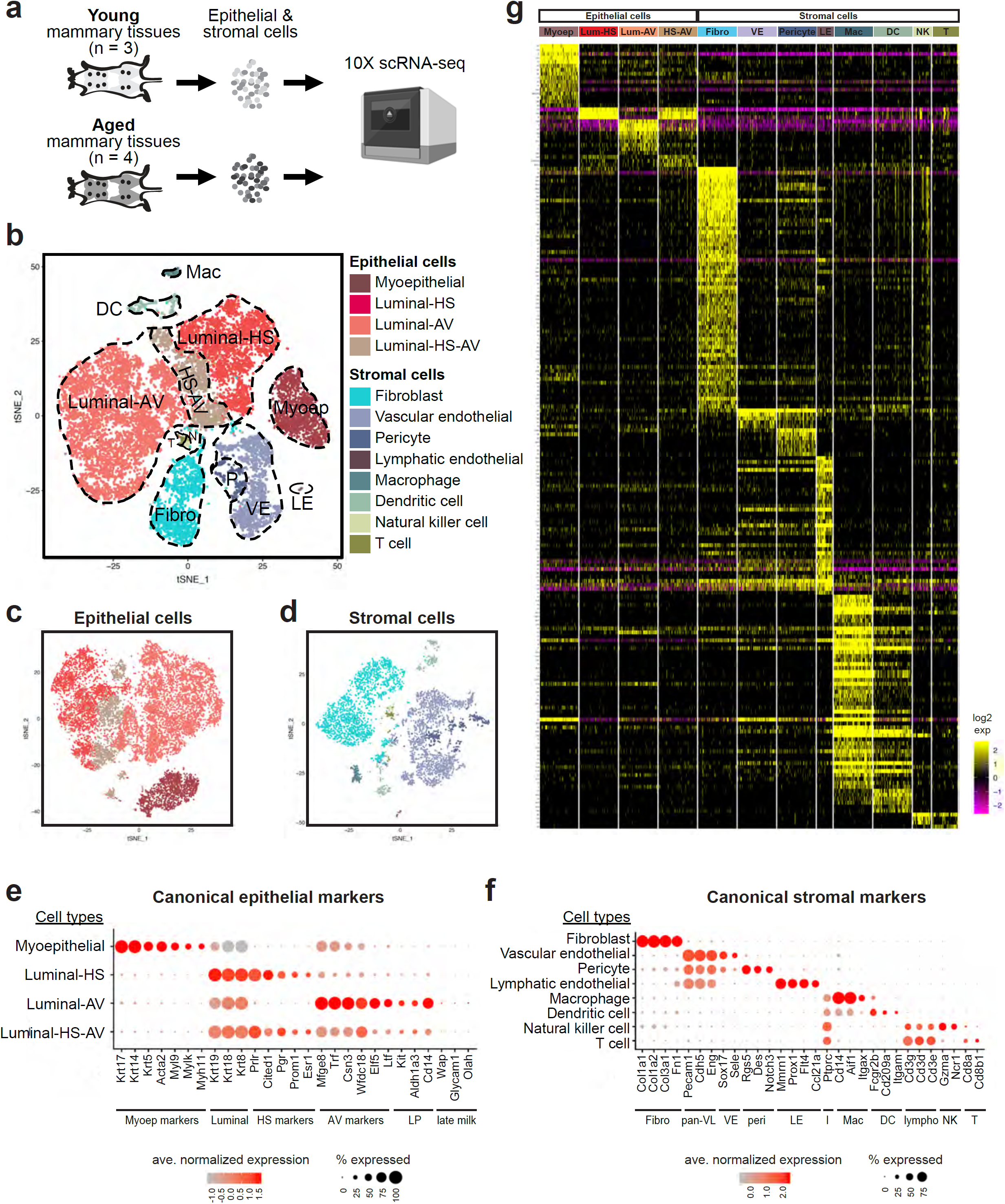
scRNA-seq analysis identifies a diversity of epithelial and stromal cells in young and aged mammary glands. **a** Overview of analysis approach. **b** t-SNE plot showing clusters of single cells. Epithelial and stromal cell types were identified based on their expression of characteristic gene markers. **c-d** Epithelial cells (**c**) and stromal cells (**d**) were subsetted and re-analyzed separately to show their respective clustering patterns. **e** Expression of known canonical markers for epithelial cell types. **f** Expression of known canonical markers for stromal cell types. **g** Gene expression heatmap of cell type-specific signatures. Abbreviations: Myoep, myoepithelial cells; HS, hormone-sensing; AV, alveolar; LP, luminal progenitors; Fibro, fibroblasts; pan-VL, pan-vascular/lymphatic cells; VE, vascular endothelial cells; LE, lymphatic endothelial cells; I, immune cells; Mac, macrophages; DC, dendritic cells; lympho, lymphocytes; NK, natural killer cells; T, T cells.

Among the epithelial cells (n = 13,939, expressing *Epcam*), we identified the two major cell types within the mammary gland – myoepithelial cells (n = 1,791, expressing *Krt17, Krt14, Krt5, Acta2, Myl9, Mylk*, and *Myh11*) and luminal cells (n = 12,148, expressing *Krt19, Krt18*, and *Krt8*). Within the luminal population, we further identified the HS cells (n = 3,571, expressing *Prlr, Cited1, Pgr, Prom1*, and *Esr1*) and AV cells (n = 6,959, expressing *Mfge8, Trf, Csn3, Wfdc18, Elf5*, and *Ltf*) (Fig. 1b, c, and e). These AV cells were in an alveolar progenitor state, as reflected by their high expression of progenitor markers, e.g. *Kit, Aldh1a3*, and *Cd14* (Fig. 1e). Furthermore, they lacked expression of late-activating milk genes^2^, e.g. *Wap, Glycam1*, and *Olah* (Fig. 1e), which is consistent with the fact that these mammary glands were not in active gestation or lactation. Nonetheless, their expression of other milk-related genes (e.g. *Mfge8, Trf, Csn3, Wfdc18*, and *Ltf*) signifies their alveolar differentiation potential in virgin non-lactating glands^15^. In addition to the HS and AV cells, we identified a subset of luminal cells that was less well characterized – a population co-expressing both hormone-sensing and alveolar genes (n = 1,618) (Fig. 1b, c, e, and g). We termed these HS-AV cells. These cells also expressed markers of luminal progenitors (e.g. *Kit, Aldh1a3*, and *Cd14*) (Fig. 1e). This population is unlikely to be a doublet artifact, as evidenced by its low doublet score (Supplementary figure 1a).

Among the stromal cells (n = 5,222), we identified a wide variety of cell types, including fibroblasts (n = 2,135, expressing *Col1a1, Col1a2, Col3a1*, and *Fn1*), vascular/lymphatic cells (n = 2,449, expressing *Pecam1, Cdh5*, and *Eng*), and immune cells (n = 638, expressing *Ptprc*, which encodes Cd45) (Fig. 1b, d, and f). The fibroblast cluster is composed of a largely homogeneous population. The vascular/lymphatic population includes vascular endothelial cells (n = 2,010, expressing *Sox17* and *Sele*), pericytes (n = 398, expressing *Rgs5, Des*, and *Notch3*), and a small set of lymphatic endothelial cells (n = 41, expressing *Mmrn1, Prox1, Flt4*, and *Ccl21a*). Finally, the immune population contains myeloid cells, namely macrophages (n = 149, expressing *Cd14, Aif1*, and *Itgax*) and dendritic cells (n = 375, expressing *Fcgr2b, Cd209a* and *Itgam*), as well as lymphocytes marked by *Cd3d, Cd3e*, and *Cd3g*, including natural killer (NK) cells (n = 47, expressing *Gzma* and *Ncr1*) and Cd8+ T cells (n = 53, expressing *Cd8a* and *Cd8b1*). We also detected very small sets of Cd4+ T cells (n = 8, expressing *Cd4* but not *Cd8a* or *Cd8b1*) and B cells (n = 6, expressing *Blnk, Cd79a*, and *Cd79b*), but their extremely low abundance prevented us from further analysis of these cell types.

Of note, the ability to capture relatively rare populations such as lymphatic endothelial cells and different immune cells, as well as the ability to detect unanticipated populations such as the HS-AV cells, highlight the power of scRNA-seq to analyze diverse cell types in the mammary gland in a more comprehensive manner without *a priori* knowledge required to pre-isolate these populations. The only major population not captured in our analysis was adipocytes, which were removed with the supernatant during the tissue dissociation process. Alternative dissociation protocols optimized for adipocyte isolation^16,17^ will be better suited to analyze these cells.

We generated gene signatures characterizing each cell type by multiple pair-wise differential gene expression analyses between a particular cell type and the other cell populations (see Methods). The resulting cell type-specific gene signatures are summarized in Fig. 1g and Supplementary table 1. These signatures can serve as a useful resource for future studies to identify or isolate a specific cell type within a heterogeneous population.

### Prevalent alterations in cell proportions and gene expression in aged mammary glands

Our scRNA-seq analysis revealed differences between young and aged mammary glands both in terms of cell type composition and gene expression pattern. An overview of these results is summarized in Figure 2 and Supplementary figure 2. For cell type composition, the relative contribution of each cell type is well conserved across samples within each age group, but differs between young versus aged mammary glands (Fig. 2a-d, Supplementary figure 1b, and Supplementary tables 2 and 3). The proportion of epithelial cells relative to stromal cells increased significantly with age, with epithelial cells expanding from 57% of total cells in young mammary glands to 85% in age tissues, while stromal cells decreased commensurately from 43% to only 15% (Fig. 2a; *p* < 0.0001, Fisher’s exact test). Age-associated changes in cell type composition within the epithelial and stromal compartments also occurred. Within the epithelial compartment, the most striking change occurred in the luminal AV cells, which exhibited a 4-fold increase in aged mammary glands compared to young tissues (Fig. 2b; *p* < 0.0001, Fisher’s exact test). This increase in luminal AV cells was accompanied by a commensurate 3-fold decrease in luminal HS cells in aged mammary epithelia (Fig. 2b; *p* < 0.0001, Fisher’s exact test). Within the stromal compartment, the most prominent change occurred in the fibroblasts, with a 4-fold decrease in aged mammary stromal cells compared to their young counterparts (Fig. 2c; *p* < 0.0001, Fisher’s exact test). The relative proportion of total immune cells decreased by 2-fold with age within the stromal compartment (Fig. 2c; *p* < 0.0001, Fisher’s exact test). Furthermore, the composition of certain immune cell types was altered with aging. The relative proportion of dendritic cells decreased by 4-fold within total immune cells in aged mammary glands compared to young tissues (Fig. 2d; *p* < 0.0001, Fisher’s exact test), whereas the relative proportion of T cells increased by 9-fold (Fig. 2d; *p* < 0.0001, Fisher’s exact test).

**Fig. 2.**
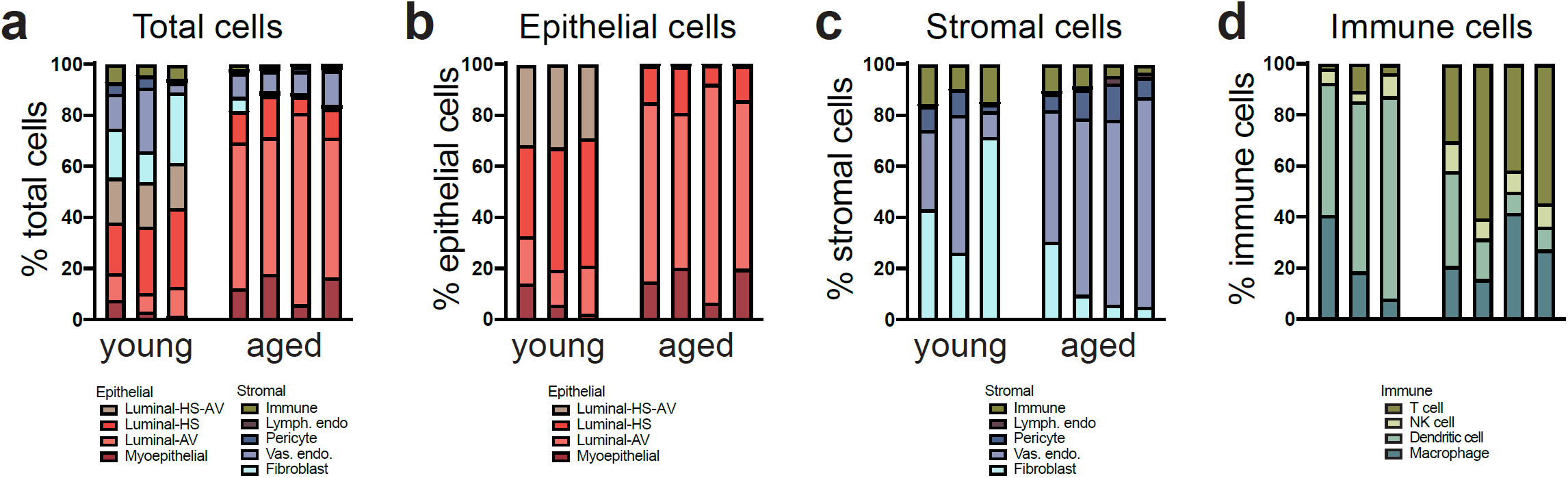
Overview of cell type composition in young and aged mammary glands. **a**-**d** Relative proportions of specified cell types among (**a**) total cells, (**b**) epithelial cells, (**c**) stromal cells, and (**d**) immune cells.

In addition to cell type composition, cell intrinsic gene expression differences also exist between young and aged mammary glands within each epithelial and stromal cell type (Supplementary figure 2a-l). Within the epithelial compartment, myoepithelial cells exhibited the largest number of differentially expressed genes (Supplementary figure 2a), whereas within the stromal compartment, vascular endothelial cells showed the most striking pattern of differential gene expression (Supplementary figure 2f). These age-associated changes are examined in greater detail below for each cell type. Overall, our results indicate that aging induces a large number of alterations both at the cell composition level and gene expression level.

### Myoepithelial cells altered in gene expression

We detected myoepithelial cells in both young and aged mammary glands. As mentioned above, these cells expressed canonical myoepithelial markers, such as *Krt17, Krt14*, and *Krt5* (Fig. 3a). The proportion of myoepithelial cells remained relatively constant with age (Fig. 3b and Supplementary table 3). However, differential gene expression analysis (see Methods) identified a large number of significant transcriptional alterations across all aged samples compared to young samples (57 genes upregulated and 72 genes downregulated with age; Fig. 3c and Supplementary table 4). Pathway analysis indicated that these differentially expressed genes primarily fell into four types of gene signatures – paracrine/juxtacrine signaling ligands, oxidative phosphorylation, extracellular matrix (ECM), and cytoskeletal genes (Fig. 3d). Specifically, aged myoepithelial cells were characterized by increased expression of multiple cytokines/chemokines that are also known to be expressed in immune cells (including *Cxcl1, Cxcl2, Cxcl16, Csf1*, and *Csf3*) as well as the juxtacrine ligand *Jag1* (Fig. 3e). Furthermore, aged myoepithelial cells exhibited reduced expression of oxidative phosphorylation genes (*Ndufa3, Ndufa13, Ndufb10, Ndufc1, Atp5j*, and *Etfb*) (Fig. 3f), which may reflect their reduced metabolic activity. Finally, these cells showed a marked reduction in the expression of many ECM-related genes (*Dcn, Col4a1, Col4a2, Emid1, Spon2*, and *Sparc*) (Fig. 3g) and multiple cytoskeletal genes (*Krt15, Acta2, Actg2, Mylk, Myl9*, and *Myh11*) (Fig. 3h), indicating a potential impairment in their functions to synthesize basement membrane proteins and maintain contractility as they age.

**Fig. 3.**
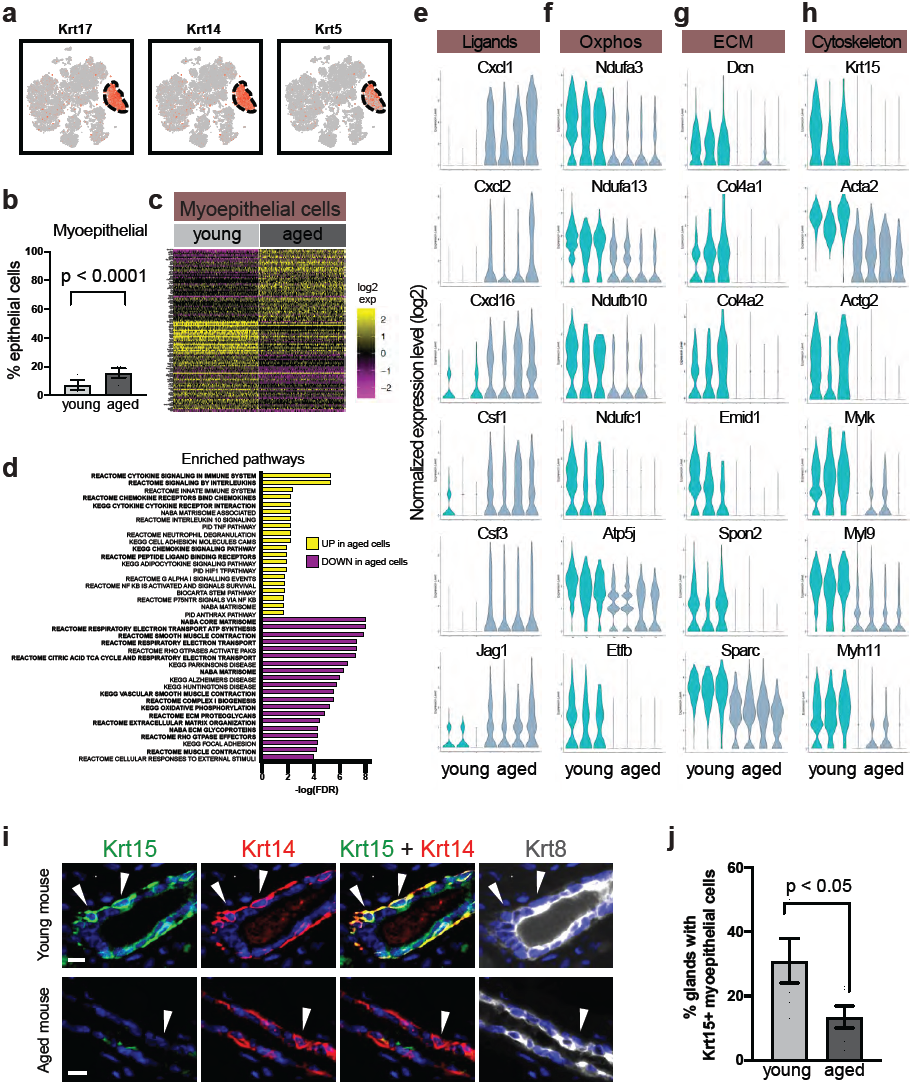
Aged myoepithelial cells show altered gene expression. **a** Myoepithelial cells were identified by their expression of characteristic gene markers in scRNA-seq analysis. **b** Relative proportion of myoepithelial cells in young (n = 3) and aged (n = 4) mammary glands as detected by scRNA-seq (Fisher’s exact test). **c** Differentially expressed genes in aged versus young myoepithelial cells. Gene names are also listed in Supplementary table 4. **d** Top gene sets identified by pathway enrichment analysis of differentially expressed genes in **c**, with recurring gene sets highlighted in bold. **e-h** Violin plots showing expression of select genes from **d** in young and aged myoepithelial cells. **i** Representative immunofluorescence staining pattern of Krt15 in myoepithelial cells (stained as Krt14+ and Krt8-, as indicated by arrow heads) in young and aged mammary glands. Scale bar = 10 *µ*m. **j** Quantification of Krt15+ myoepithelial cells in **i** (n = 6 animals per age group, Student’s t test).

Of note, because *Krt15* has been reported as a myoepithelial marker in murine mammary glands^1,2^, the reduced expression of *Krt15* (3.8-fold decrease, adjusted *p*-value < 10^−108^, Wilcoxon rank sum test) in aged mammary glands would suggest that it may not be a reliable marker for aged myoepithelial cells. We therefore performed immunofluorescence staining to confirm Krt15 protein expression pattern in young and aged myoepithelial cells, using Krt14 as a reference myoepithelial marker as its expression level did not change with age as detected by scRNA-seq (fold difference = 1.0, adjusted *p*-value = n.s., Wilcoxon rank sum test). Consistent with the scRNA-seq data, immunofluorescence staining demonstrated that Krt15 was expressed in Krt14+ myoepithelial cells in young mammary glands, but its expression was significantly reduced in aged mammary glands (Fig. 3i and 3j). These expression patterns suggest that Krt15 should be used with caution as a myoepithelial marker depending on the age of the mice.

### Luminal HS and AV cell changed in proportions

ScRNA-seq captured two canonical luminal populations – HS cells and AV cells. These cells were identified by their characteristic markers, namely, hormone-sensing genes *Prlr, Pgr*, and *Esr1* for HS cells (Fig. 4a) and milk-related genes *Mfge8, Ltf*, and *Csn3* for AV cells (Fig. 4b). Interestingly, these populations exhibited striking changes in their relative proportion with age. Whereas HS cells accounted for 45% of epithelial cells in young mammary glands, their representation decreased to only 14% in aged tissues (Fig. 4c). On the other hand, AV cells showed a commensurate increase from 17% of epithelial cells in young mammary glands to 71% in aged tissues (Fig. 4d).

**Fig. 4.**
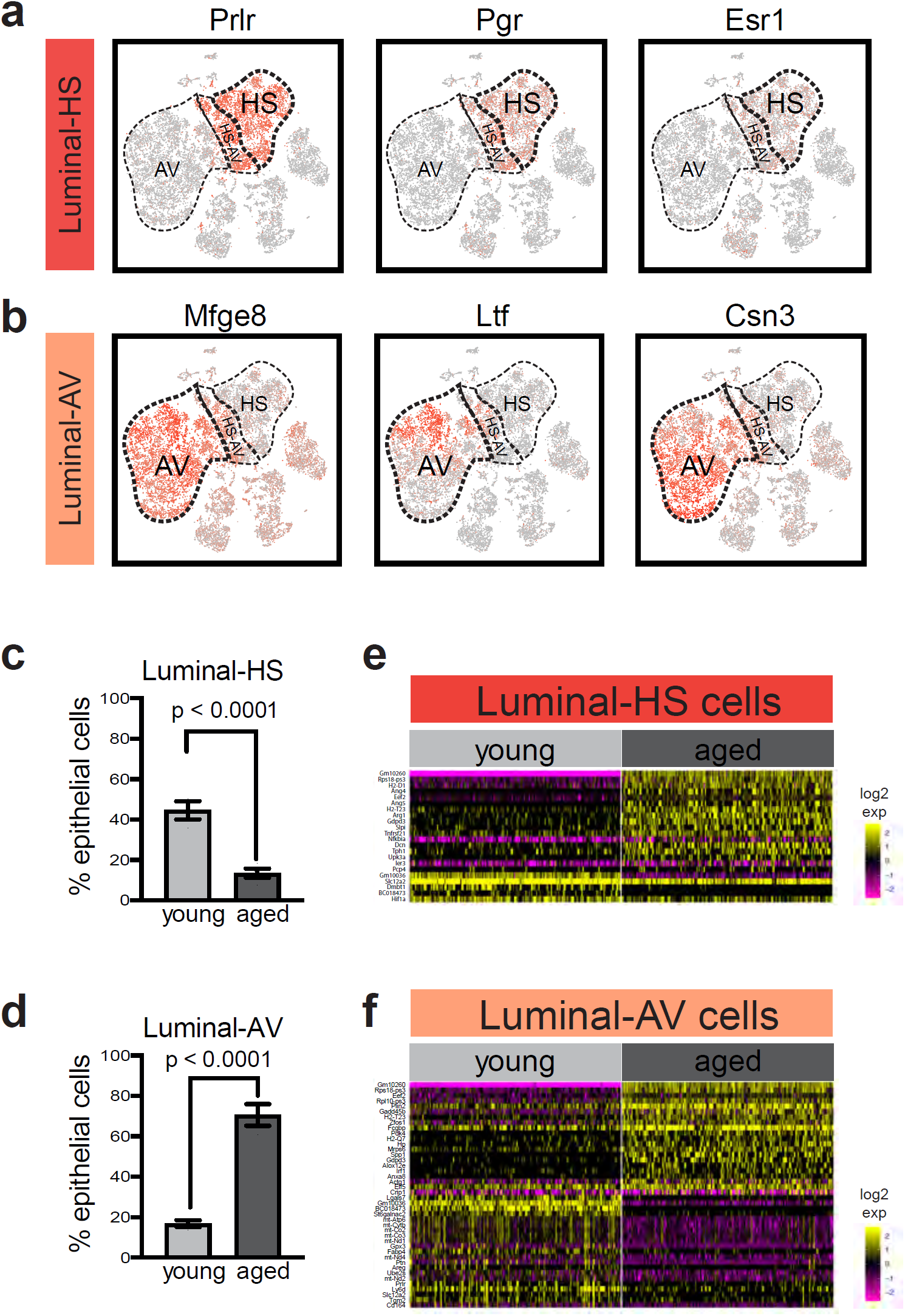
Alterations in aged luminal hormone-sensing (HS) cells and alveolar (AV) cells. **a** HS cells are marked by the expression of hormone receptors in scRNA-seq analysis. **b** AV cells are marked by the expression of milk-related genes. **c** Relative proportion of HS cells in young (n = 3) and aged (n = 4) mammary glands (Fisher’s exact test). **d** Relative proportion of AV cells in young (n = 3) and aged (n = 4) mammary glands (Fisher’s exact test). **e** Differentially expressed genes in young versus aged HS cells. **f** Differentially expressed genes in young versus aged AV cells. For **e** and **f**, gene names are also listed in Supplementary table 4.

Differential gene expression analysis revealed a relatively small number of aging-associated transcriptional changes in HS and AV cells compared to myoepithelial cells. In aged HS cells, 17 genes were significantly increased in expression (with the top genes being *Dcn, Slpi, Upk3a, Ang4*, and *Ang5*) and five genes showed decreased expression (*Dmbt1, Slc12a2, Hif1a, Gm10036*, and *BC018473*) compared to young HS cells (Fig. 4e and Supplementary table 4). Interestingly, *Dmbt1, Slc12a2*, and *Hif1a* all have known roles in mammary development or disease, and are preferentially expressed in HS cells instead of AV cells in normal mammary glands. *Dmbt1* is a putative tumor suppressor gene that is downregulated in breast cancer relative to normal mammary tissues^18,19^, whereas *Slc12a2* (also known as *Nkcc1* or *Bsc2*, encoding the Na-K-Cl cotransporter) and *Hif1a* (a component of a hypoxia-responsive master transcription factor) are required for normal epithelial differentiation during mammary development^20,21^. The effects of their downregulated expression in aged HS cells warrants further investigations in the future.

Similar gene expression analysis in AV cells identified 21 upregulated genes (top examples included *Spp1, Hp, Gadd45b, Fcgbp, Alox12e*, and *Plin2*) and 21 downregulated genes (top examples included *Ptn, Gpx3, Lgals7*, and *Fabp4*) in aged compared to young tissues (Fig. 4f and Supplementary table 4). Of note, the decrease in expression of the redox regulators *Gpx3* and *Fabp4* in aged AV cells may reflect a change in their redox state. In addition, the enhanced expression of the master alveolar-lineage transcription factor *Elf5* and milk-related genes *Spp1*^22-24^ and *Plin2*^25,26^ in aged AV cells may potentially reflect their shift toward a slightly more mature state in the alveolar lineage, consistent with previous report^27^ as well as our observation (Supplementary figure 3) that aged mammary glands showed signs of activated alveolar cells and increased secretory material independent of pregnancy.

### Luminal HS-AV cells showed age-dependent abundance

As mentioned above, our scRNA-seq analysis also suggested the presence of another luminal cell subset – HS-AV cells that co-expressed both luminal HS and AV markers as well as luminal progenitor markers (Fig. 1e, 4a-b). While similar cells were captured in two recent scRNA-seq analyses of mouse mammary epithelial cells^1,2^, they have not been characterized further. In addition to expressing both luminal HS and AV markers, HS-AV cells are also enriched for other markers when compared to HS cells and AV cells. We performed gene expression analysis to identify markers that distinguished the HS-AV population from HS cells and AV cells, and we performed the analysis within the young mammary gland in order to focus on cell type-specific difference without the effects of aging. The results revealed that HS-AV cells were enriched for 15 genes and depleted for 5 genes when compared to luminal HS cells or AV cells (Fig. 5a and Supplementary table 5). Among the enriched genes were three transcription factors, *Sox9, Bhlhe41*, and *Nfia* (Supplementary figure 4a-c). While *Sox9* has been implicated in promoting a stem/progenitor state in mammary cells^28^, the roles of *Bhlhe41* (also known as *Sharp1* or *Dec2*) and *Nfia* in normal mammary development are not well understood and warrant additional studies in the future. Moreover, the progesterone-dependent acylglycerol lipase *Abhd2* is highly enriched in HS-AV cells when compared to HS cells (2.2-fold, *p* < 10^−23^) or AV cells (3.3-fold, *p* < 10^−110^) (Supplementary figure 4d and Supplementary table 5). However, the role of *Abhd2* in mammary gland biology is not known and deserves future investigations.

**Fig. 5.**
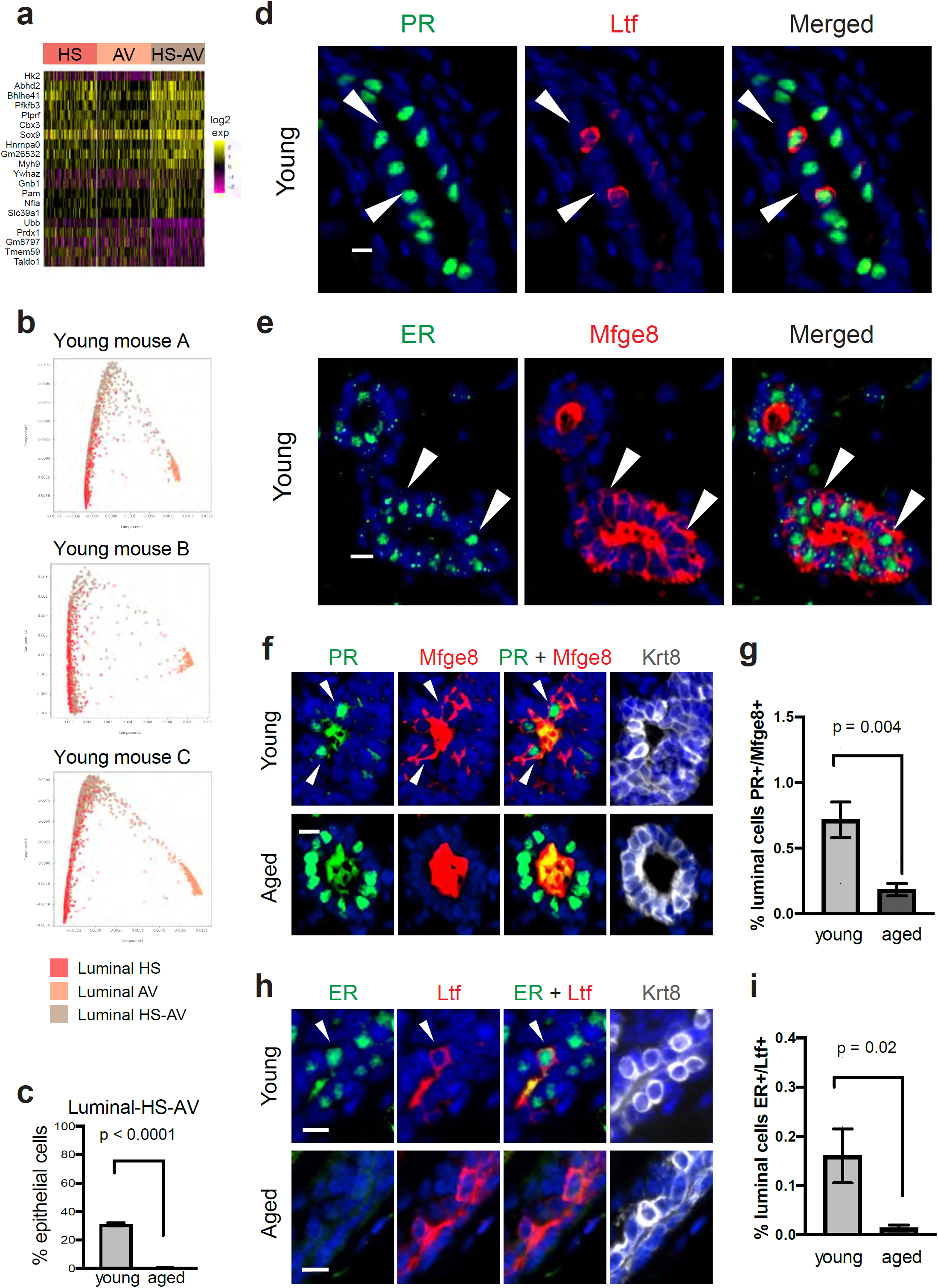
Detection of HS-AV luminal cells in young and aged mammary glands. **a** Gene markers distinguishing HS-AV cells from both HS cells and AV cells in scRNA-seq analysis of young mammary glands. **b** Dimensionality reduction plots from STREAM lineage trajectory analysis of luminal cells in young mammary glands (n = 3 samples). **c** Relative proportion of HS-AV cells in young (n = 3) and aged (n = 4) mammary glands as detected by scRNA-seq (Fisher’s exact test). **d** and **e** Representative immunofluorescence staining pattern of co-expression of HS marker (PR or ER) and AV marker (Ltf or Mfge8) in HS-AV cells of young mammary glands. **f** and **g** Immunofluorescence staining to identify (**f**) and quantify (**g**) HS-AV cells co-expressing PR and Mfge8 in young and aged mammary glands. **h** and **i** Immunofluorescence staining to identify (**h**) and quantify (**i**) HS-AV cells co-expressing ER and Ltf in young and aged mammary glands. In **g** and **i**, n = 6 animals per age group were analyzed by Student’s t-test. In all immunofluorescence staining panels, scale bar = 10 *µ*m. Mfge8 and Ltf proteins were also detected as secreted proteins in the lumen of the mammary glands. In **f** and **h**, Krt8 marks luminal cells.

To examine the lineage relationship between HS-AV cells to HS cells and AV cells, we performed STREAM lineage trajectory analysis within each of the three young mammary glands analyzed by scRNA-seq^29^. HS-AV cells primarily localized to the bifurcation junction between HS cells and AV cells (Fig. 5b and Supplementary figure 5). The pattern is highly reproducible across all three young samples. This predicted lineage trajectory of the HS-AV cells relative to HS cells and AV cells is consistent with their gene expression pattern being a hybrid between these two luminal cell types, and suggests the possibility that HS-AV cells might have the potential to differentiate into the HS and AV lineages.

To confirm the existence of HS-AV cells in mammary tissues, we performed *in situ* immunofluorescence staining in a panel of young mammary glands (n = 6) using established hormone-sensing markers, namely Progesterone Receptor (PR) and Estrogen Receptor (ER), and alveolar markers, namely Lactotransferrin (Ltf) and Milk Fat Globule-EGF Factor 8 (Mfge8). This analysis confirmed the presence of HS-AV cells co-expressing PR/ER with Ltf/Mfge8 (Fig. 5d-e). HS-AV cells were localized in a scattered pattern within both ductal and alveolar regions of the mammary glands. PR+/Mfge8+ cells and ER+/Ltf+ cells were present at 0.72% and 0.16% of luminal cells, respectively (Fig. 5f-i). Of note, the lower abundance of ER+/Ltf+ cells compared to PR+/Mfge8+ cells by immunofluorescence staining is consistent with the lower abundance of Ltf+ cells compared to Mfge8+ cells in the HS-AV population in scRNA-seq (Fig. 4b). Furthermore, while we were able to detect HS-AV cells both by scRNA-seq and immunofluorescence staining, we noted that these cells were lower in abundance when analyzed at the protein level by immunofluorescence than at the RNA level by scRNA-seq (Fig. 5c, g, and i). This difference may reflect gene expression regulations at the post-transcriptional and post-translational levels.

Interestingly, the abundance of these HS-AV cells diminished dramatically with age. In our scRNA-seq analysis, HS-AV cells accounted for an average of 31% of epithelial cells in young mammary glands but only 0.4% in aged mice, an 82-fold decrease in relative proportion (Fig. 5c). We confirmed this pattern by immunofluorescence staining of young and aged mammary glands (n = 6 per group) using a combination of HS and AV markers, and found that HS-AV cells indeed decreased when we stained for PR+/Mfge8+ luminal cells (Fig. 5f-g, 4-fold decrease). Additional staining with a second HS and AV marker combination, ER and Ltf, further confirmed this decrease in HS-AV cell abundance (Fig. 5h-i, 12-fold decrease). Collectively, these results demonstrated an age-dependent existence of the HS-AV cells.

### Stromal cells changed in both proportions and gene expression

In addition to epithelial cells, our analysis captured a variety of stromal cells in both young and aged mammary glands (Fig. 6a-d). The most abundant stromal cell type was fibroblasts (Fig. 6a). These cells homogeneously expressed ECM genes (e.g. *Fn1, Col1a1, Col1a2, Col3a1*) but not contractile genes (e.g. *Acta2, Myl9, Mylk, Myh11*) (Fig. 6a and Supplementary figure 6), indicating that they were ECM-producing fibroblasts instead of contractile myofibroblasts. Interestingly, aged fibroblasts exhibited both cell-level and gene-level alterations. The relative proportion of fibroblasts decreased from 47% of stromal cells in young mammary glands to only 13% in aged mammary glands (Fig. 6e). In addition, differential gene expression analysis revealed 15 genes upregulated and 16 genes downregulated with age. The vast majority of aged fibroblasts gained expression of several stress-related genes (*Hspa1a, Gadd45b*, and *Cebpb*), while losing expression of several ECM-related genes (*Fn1, Col6a3*, and *Mmp23*) (Fig. 6i-k and Supplementary table 4). Notably, *Fn1*, which encodes fibronectin, is a major component of the ECM and a master organizer of matrix assembly^30-32^. The decrease in both fibroblast proportion and ECM gene expression is consistent with previous reports in human mammary glands showing that aging is associated with a reduction of connective tissues^33,34^.

**Fig. 6.**
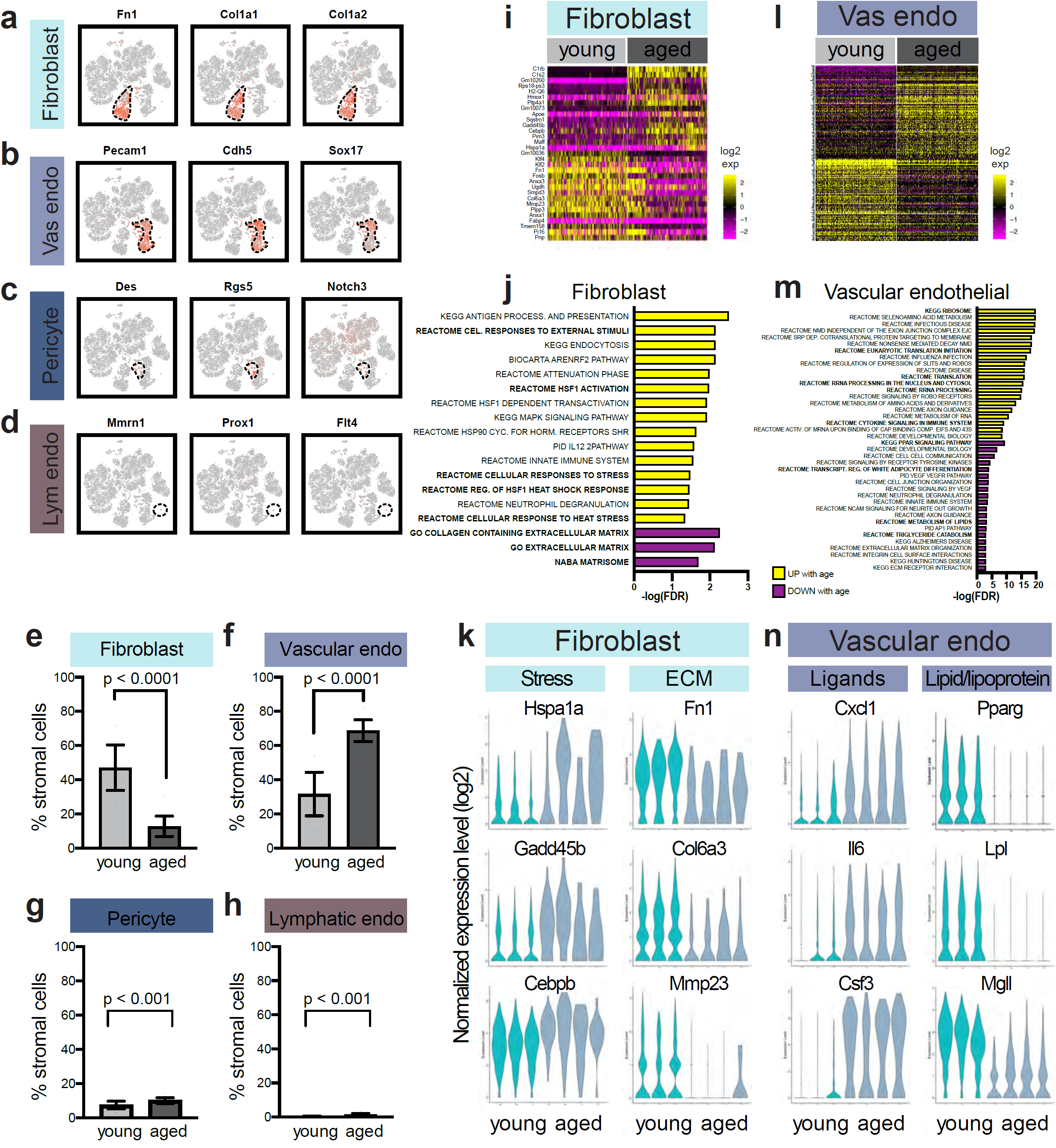
Age-dependent alterations in stromal fibroblasts and vascular/lymphatic cells. **a**-**d** Fibroblasts, vascular endothelial cells, pericytes, and lymphatic endothelial cells are distinguished by their expression of respective characteristic gene makers in scRNA-seq. **e**-**h** Relative proportions of fibroblasts, vascular endothelial cells, pericytes, and lymphatic endothelial cells in young (n = 3) and aged (n = 4) mammary glands in scRNA-seq analysis (Fisher’s exact test). **i** Differentially expressed genes in aged versus young fibroblasts. **j** Top gene sets identified by pathway enrichment analysis of differentially expressed genes in **i**, with recurring gene sets highlighted in bold. **k** Violin plots of select genes identified in **i** and **j** for fibroblasts. **l** Differentially expressed genes in aged versus young vascular endothelial cells. **m** Top gene sets identified by pathway enrichment analysis of differentially expressed genes in **l**, with recurring gene sets in bold. **n** Violin plots of select genes identified in **l** and **m** for vascular endothelial. For **i** and **l**, gene names are also listed in Supplementary table 4.

We also detected vascular/lymphatic cells, including vascular endothelial cells (expressing *Pecam1, Chd5*, and *Sox17*), pericytes (expressing *Des, Rgs5*, and *Notch3*), and lymphatic endothelial cells (expressing *Mmrn1, Prox1*, and *Flt4*) (Fig. 6b-d). Among these cells, vascular endothelial cells exhibited the most dramatic changes in terms of cell proportion and gene expression (Fig. 6f and l). Vascular endothelial cells increased in proportion from an average of 32% of stromal cells in young mammary glands to 69% in aged tissues (Fig. 6f). In addition, aged vascular endothelial cells exhibited 102 genes upregulated and 90 genes downregulated with age (Fig. 6l and Supplementary table 4). Pathway analysis revealed that the signatures of upregulated genes were enriched for cytokines/chemokines (*Cxcl1, Il6*, and *Csf3*), whereas the downregulated genes were enriched for lipid/lipoprotein metabolism (e.g. *Pparg, Lpl, Mgll, Fabp4, Cd36*, and *Plin2*) (Fig. 6m-n and Supplementary table 4). In contrast, the proportion of pericytes and lymphatic endothelial cells within the stroma remained relatively constant with age, and their gene expression also remained largely unaltered between young and aged mammary glands (Fig. 6g-h and Supplementary figure 2).

Finally, our scRNA-seq data captured different types of immune cells in both young and aged mammary glands. These included myeloid cells (macrophages marked by the expression of *Cd14, Aif1*, and *Itgax*; and dendritic cells marked by the expression of *Fcgr2b, Cd209a*, and *Itgam*) as well as lymphocytes (NK cells marked by the expression of *Gzma, Ncr1*, and *Cd244*; and T cells marked by the expression of *Cd8a, Cd8b1*, and *Il7r*) (Fig. 7a-d). These immune cells represented tissue-resident immune cells, as we resected and discarded the lymph nodes in our mammary sample preparation prior to scRNA-seq. Interestingly, we observed that the proportion of dendritic cells and T cells changed with age (Fig. 7e-h), with myeloid dendritic cells decreasing from 66% of total immune cells to 18%, and T lymphocytes increasing drastically from 5% to 47%. In contrast, the relative abundance of macrophages and NK cells remained relatively constant across young and aged mammary glands. In terms of gene expression changes, macrophages showed a more prominent alteration pattern compared to the other immune cell types (11 genes upregulated and 9 genes downregulated with age; Fig. 7i and Supplementary table 4). Pathway analysis of the genes upregulated in aged macrophages uncovered an enrichment of chemokines (e.g. *Ccl4, Ccl5*, and *Cxcl2*) (Fig 7j-k). Notably, *Ccl4* and *Ccl5* are T-cell recruiting chemokines^35^, and their increased expression is consistent with the higher infiltration of T cells observed in the aged mammary glands.

**Fig. 7.**
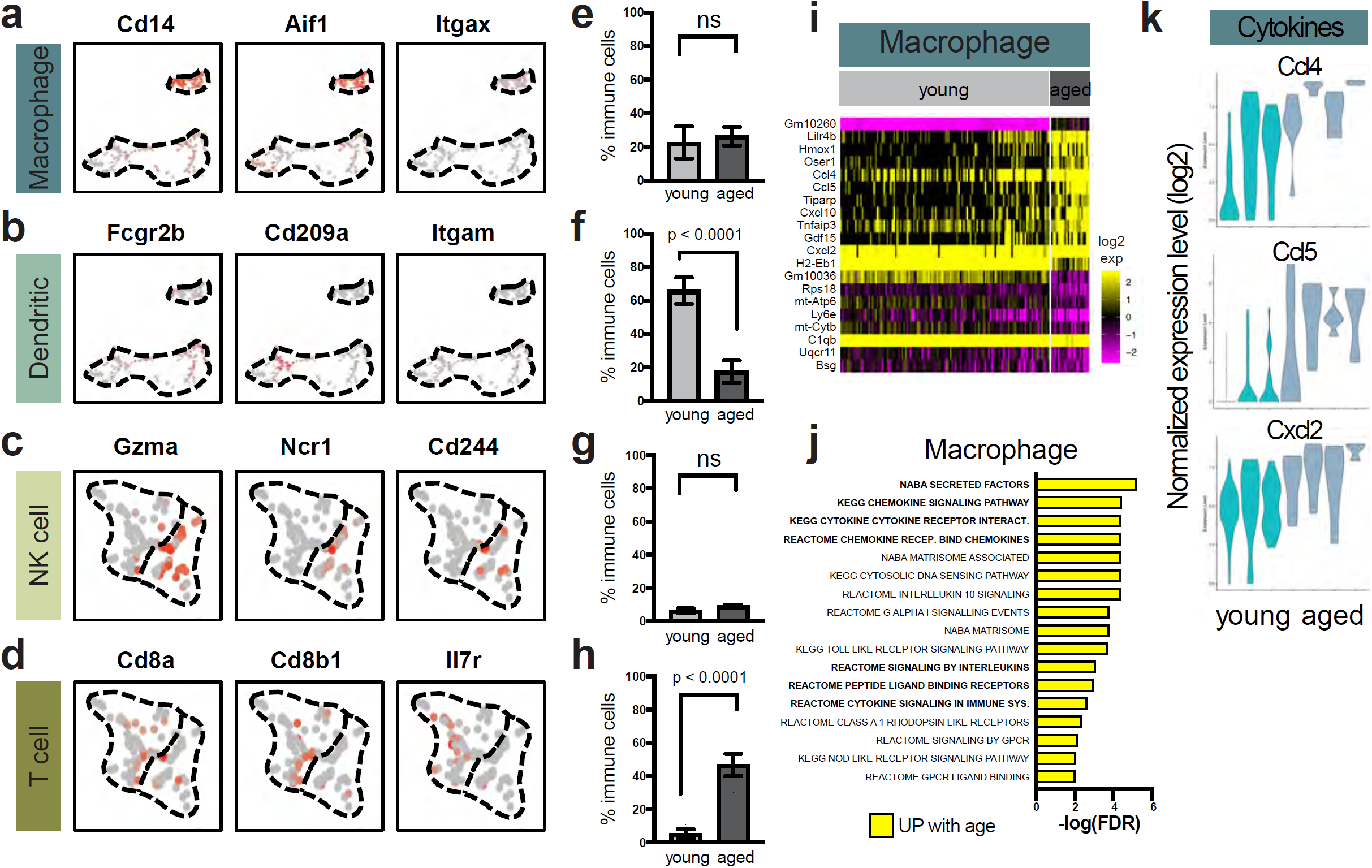
Immune cells exhibit alterations in young and aged mammary glands. **a**-**d** Macrophages, dendritic cells, natural killer (NK) cells, and T cells are distinguished by their expression of respective canonical gene markers in scRNA-seq. **e**-**h** Relative proportions of macrophages, dendritic cells, NK cells, and T cells in young (n = 3) and aged (n = 4) mammary glands as detected by scRNA-seq (Fisher’s exact test). **i** Differentially expressed genes in aged versus young macrophages. **j** Top gene sets identified by pathway enrichment analysis of differentially expressed genes in **i**, with recurring gene sets highlighted in bold. Gene names are also listed in Supplementary table 4. **k** Violin plots of select genes identified in **i** and **j** for macrophages.

## Discussion

The single-cell transcriptomic profiles shown here capture the age-associated alterations in the mammary epithelium and stroma at high resolution, revealing numerous changes at both the cell and gene levels. The young and aged murine samples analyzed correspond to human mammary glands in early adulthood (20-30 years of age) and perimenopause (45-55 years of age), respectively^36^. Considering the longer lifespan of humans compared to mice, there may be additional age-related alterations in the human mammary glands that are not captured by a murine model. However, unlike human tissues, mouse mammary glands allow for the assessment of the alterations associated with intrinsic physiological changes directly related to aging, without the influence of exogenous environmentally-induced stresses unavoidable in human studies, such as diet, alcohol consumption, and exposure to carcinogens, as well as history of pregnancy and lactation. Importantly, the cell- and gene-level profiles of the mammary gland are consistent within each age group to a striking degree, suggesting that the intrinsic effects of aging are well conserved. Therefore, the data described herein complement previous studies on pregnancy-related mammary gland changes, and provide a useful resource for better understanding the intrinsic effects of aging on the mammary gland.

Aging induces shifts in the biological functions of various mammary cells. For example, within the mammary epithelium, the dramatic expansion of the AV population with age, from a low-abundance cell type in young mammary glands to the most dominant population in older glands, reflects a significant increase in proliferation of these cells relative to the other cell types. In addition to their increase in relative proportion, aged AV cells also exhibit higher expression levels of genes associated with mature alveolar cells, including the master alveolar transcription factor *Elf5*^37,38^, as well as milk-related genes *Plin2*^25,26^ and *Spp1*^22-24^, suggesting a shift toward a more differentiated mature alveolar state with age. These changes are consistent with previous report^27^ as well as our observation that aged nulliparous mice exhibit higher levels of budding alveoli and dilated ducts filled with secretory material without pregnancy, and are likely caused by the continuous stimulation by hormones in the estrous cycle^27^. Age-induced alterations also occur in myoepithelial cells. Two major functions of myoepithelial cells in the mammary gland are to synthesize basement membrane proteins^39,40^ and to mediate contraction to disperse milk. Aged myoepithelial cells exhibit diminished expression of several type IV collagens as well as cytoskeletal genes, which could reflect an age-associated decline in their normal functions in maintaining the basement membrane and contractility. In addition to the decreased ECM synthesis by myoepithelial cells, fibronectin and type VI collagen production by fibroblasts is also diminished at the transcription level and the relative proportion of fibroblasts was reduced. Together, these changes could contribute to the overall loss of ECM deposition and tissue density in aged mammary glands^33,34,41,42^. Finally, myoepithelial cells, vascular endothelial cells, and macrophages exhibit increased expression of several inflammatory cytokines and chemokines, including *Cxcl1, Cxcl2, Ccl4, Ccl5*, and *Csf3*, which may influence the infiltration and function of immune cells such as T lymphocytes and therefore indirectly affect mammary epithelial cells via stromal-epithelial interactions. Overall, these findings highlight the diversity and complexity of aging-induced alterations within the mammary gland.

Given that the aged mammary gland is associated with increased risk of diseases such as cancer both in mice and in human^27,43^, our findings may also provide initial insights into the underlying mechanisms of how aging contributes tumor development. For instance, alveolar luminal progenitor cells have been hypothesized as the cells-of-origin for triple-negative breast cancer^44-46^. The expansion of AV luminal progenitor cells with age may provide a mechanism for the increased breast cancer risk in older women. Moreover, myoepithelial cells have been demonstrated to confer a tumor suppressive function by collectively acting as a dynamic contractile barrier to restrain and reinternalize migrating invasive luminal cells^47^. This function is dependent on the expression of α-smooth muscle actin, the product of the gene *Acta2* ^47^. The decreased expression of *Acta2* and other contractile cytoskeletal genes in aged myoepithelial cells, together with a partially compromised basement membrane due to reduced expression of its components, may impair their ability to maintain the tumor suppressive barrier function, thereby increasing the risk of malignant outgrowth and dissemination.

The identification of HS-AV cells co-expressing HS markers and AV markers is of considerable interest. Examination of publicly available single-cell data revealed that these cells were also captured in two recent scRNA-seq studies on young nulliparous murine mammary glands^1,2^. While Bach et al. did not specifically comment on the hybrid nature of their gene expression pattern, Pal et al. described them as luminal intermediate cells with gene expression at intermediate levels between the AV progenitor lineage (marked by *Elf5, Aldh1a3, Cd14, Csn3*, and *Trf*) and the HS lineage (marked by *Prlr* and *Cited1*). Here, we confirmed the existence of cells that co-express both HS and AV proteins in the mammary gland by immunofluorescence staining, and further characterized their gene expression pattern by differential gene expression and lineage trajectory analyses. We also demonstrated that HS-AV cells decrease in abundance with age, possibly due to a number of reasons, including a lack of self-renewal, differentiation into the HS or AV lineages, or out-competition by proliferation of the HS and AV luminal cell types.

The lineage relationship of HS-AV cells to HS cells and AV cells remains to be determined. Multiple lineage tracing studies using HS lineage reporters (*Esr1* and *Prom1*) as well as AV lineage reporters (*Notch1* and *Sox9*), have demonstrated that the HS and AV lineages in the postnatal mammary glands are largely maintained by independent pools of lineage-restricted unipotent progenitor cells^48-50^. These four genes are also expressed in HS-AV cells at the transcriptional level. Given these lineage studies, it is unlikely that under normal homeostasis HS-AV cells could function as bi-potent progenitors that contribute significantly to the HS and AV lineages in the adult mammary glands. Thus, it is more likely that HS-AV cells represent a distinct and restricted population that is independent from HS cells and AV cells. Furthermore, depending on the rate of their cell division, it is possible that HS-AV cells may not form clones with sufficient size to be detectable to a significant degree in lineage tracing analyses, especially in the context of sparse labeling and short-term tracing.

If these HS-AV cells are a stable luminal population, it raises the question of what their function might be. One possibility is that they might serve as dormant resident unipotent or bipotent luminal progenitors that can become activated under tissue regeneration or repair conditions to contribute to the HS and/or AV lineages. Similar examples of dormant progenitor cells have been described in the lungs^51-53^ and liver^54,55^. Several observations provide a hint in support of this hypothesis. First, in lineage trajectory analysis based on scRNA-seq data, HS-AV cells are localized to the bifurcation of the HS branch and the AV branch, suggesting the possibility that they might be transcriptionally primed to give rise to cells in either the HS or AV lineage. Second, when compared to HS cells and AV cells, HS-AV cells are enriched for the expression of *Sox9*, a transcription factor previously reported to confer progenitor property in luminal cells^28^. Third, HS-AV cells express several luminal progenitor markers, such as *Kit, Aldh1a3*, and *Cd14*. Finally, the abundance of HS-AV cells in the mammary gland diminishes with age, consistent with the age-dependent depletion of stem/progenitor cells in certain tissue types^56-58^. Of note, the closer proximity of HS-AV cells to HS cells than to AV cells in lineage trajectory analysis would suggest the possibility that HS-AV cells might be potential unipotent progenitors of mature HS cells as opposed to AV cells; however, the increased expression of *Sox9* in HS-AV cells would suggest the possibility that they might be potential unipotent progenitors of the AV lineage, as *Sox9* has been reported to mark AV cells in a lineage tracing study using a transgenic reporter^48^. Functional differentiation and lineage tracing analyses are required to determine whether HS-AV cells exhibit any unipotent or bipotent progenitor properties that can be activated under tissue regeneration conditions. The co-expression of cell-surface HS markers (e.g. *Prom1* and *Ly6d*) and AV markers (e.g. *Kit* and *Cd14*) suggests that HS-AV cells can be isolated for further studies in *in vitro* and transplantation experiments *in vivo*. Finally, it remains possible that HS-AV cells represent a transient intermediate between the HS and AV populations as proposed by Pal et al.^1^, but evidence for such a transition has been limited and further studies will shed light on this model. Overall, the presence of HS-AV cells suggests a greater complexity of cell populations in mammary gland differentiation than previously appreciated and underscores the need for future studies in this area.

Taken together, this rich dataset of single-cell transcriptomic profiles helps elucidate the complex heterogeneity within mammary cell types, and provides a useful resource for future studies to understand the interactions between epithelial and stromal cell types within normal and diseased mammary tissues.

## Methods

### Animals

For single-cell RNA sequencing, young (3-4 months old) and aged (13-14 months old) virgin female mice of a mixed FVB, 129, C57BL/6J background were used in this study. For immunofluorescence staining, additional young (3-4 months old) and aged (16-months old) mice of the same background were used. Mice were fed on a regular diet (LabDiet 5053). Vaginal cytology was performed at the time of mammary gland collection to determine the stage of the estrous cycle, and only mice in diestrus were used to avoid confounding effects from hormonal changes. Aged mammary glands were confirmed to have normal histology by a rodent pathologist. All animal work was performed in accordance to protocols approved by the Institutional Animal Care and Use Committee at Harvard University.

### Mammary tissue dissociation for single-cell RNA sequencing

Abdominal #4 murine mammary glands, with lymph nodes removed, were finely minced and then incubated in a digestion solution containing DMEM/F12 (Gibco 11330), 10% heat-inactivated fetal bovine serum, 2 mg/ml collagenase XI (Sigma C9407), and 0.1 mg/ml hyaluronidase (Sigma H3506) for 1.5 hours at 37°C with constant shaking at 150 rpm. The dissociated cells were then subjected to red blood cell lysis (Biolegend 420301), a 5-minute treatment with 1 U/ml dispase (Stem Cell Technologies 07913) and 0.1 mg/ml DNase (Stem Cell Technologies 07900), and filtered through a 40 *µ*m cell strainer. Cells were resuspended in PBS containing 0.04% BSA, counted manually under the microscope, and adjusted for loading 7,000 viable cells for single-cell RNA sequencing.

### Single-cell RNA library preparation and sequencing

Single cell capturing and cDNA library generation were performed using the 10X Chromium 3’ library construction kit v2 following the manufacturer’s instruction. The libraries were then pooled and sequenced by Illumina HiSeqX.

### Single-cell RNA seq data processing

Paired-end reads from Illumina HiSeq were processed and mapped to the mm10 mouse genome using Cell Ranger v2.0. We filtered out cells with (1) UMI counts < 500 or > 60,000, (2) gene counts < 500 or > 6,000, and (3) mitochondrial gene ratio > 10%. This pre-filtering resulted in the detection of 17,509 genes in 19,161 cells, with approximately 1,600-4,000 cells from each sample. A median of 1,919 genes and 6,964 transcripts were captured per cell. The filtered data were then analyzed by Seurat v3 ^12,13^. Cell doublets were estimated using Scrublet^14^. Lineage trajectory analysis was performed on the luminal epithelial cells of each of the three young samples using STREAM^29^.

### Cell type-specific gene signatures

Cell type-specific gene expression signatures were identified as genes that exhibited expression levels two-fold or greater (with adjusted *p*-values < 0.05) than each of the other cell types in multiple pairwise differential expression analyses using Seurat’s FindMarkers function (Wilcoxon rank sum test). Similarly, the gene expression signature for HS-AV luminal cells was generated by identifying genes with expression levels at least 1.5-fold higher or lower (with adjusted *p*-values < 0.05) when compared to the luminal HS cells and to the luminal AV cells separately. Heatmaps for visualizing marker gene expression were median-centered and down-sampled to 100 cells per cell type.

### Differential gene expression analysis

Differential gene expression analysis was performed to compare young and aged samples within each cell type by using a combination of Seurat’s FindMarkers function (Wilcoxon rank sum test) and zingeR-edgeR zero-inflated negative binomial analysis^59^ for increased stringency. Differentially expressed genes were first identified using two criteria in FindMarkers analysis: (i) an expression difference of at least 1.5-fold and an adjusted *p*-value of < 0.05 in a grouped comparison of young mice (n = 3) versus aged mice (n = 4), with gene expression detected in at least 10% of cells in either the young or aged population; and (ii) an expression difference of at least 1.25-fold in all of the 12 possible combinations of young-versus-aged per-sample pairwise comparison. The resulting gene list from FindMarkers analysis was then further intersected with those identified by zingeR-edgeR (grouped analysis of young versus aged samples, with fold-difference cutoff at 1.5 and adjusted *p*-value cutoff at 0.05) to generate a final list of differentially expressed genes across age groups. Heatmaps for visualizing the differentially expressed genes were median-centered and down-sampled to 200 cells per age group.

### Pathway enrichment analysis

Pathway enrichment analysis was performed on differentially expressed genes using MSigDB^60,61^ curated canonical pathway database. For the rare circumstance (in the case of fibroblasts) where the canonical pathway database resulted in few enriched gene sets, a second database, GO cellular component, was also used. To ensure proper gene name mapping, all mouse gene names were converted to their human homologs using NCBI Homologene prior to analysis in MSigDB. Gene sets with FDR < 0.05 were deemed statistically significant. Up to top 20 enriched gene sets were shown.

### Histology and immunofluorescence staining

Mouse mammary tissues were fixed in 10% neutral buffered formalin overnight. Paraffin embedding and sectioning were performed by the Rodent Histopathology Core at Harvard Medical School. Immunofluorescence staining was performed using the following antibodies: Krt15 (BioLegend 833904, 1:50), Krt14 (BioLegend 905301, 1:250), Krt8 (DSHB Troma-I, 1:100; Abcam ab192467, 1:100), progesterone receptor (CST 8757, 1:25; Dako A0098, 1:20), estrogen receptor (Santa Cruz sc-542, 1:20; BioRad MCA1799T, 1:25), Ltf (Bioss bs-5810, 1:50), and Mfge8 (Thermo MA5-23913, 1:50; R&D Systems MAB2805, 1:50). All Ltf and Mfge8 antibodies have been previously validated by staining lactating mammary glands.

### Statistics

For cell type composition comparison between young and aged mammary glands in scRNA-seq, statistically significant difference was determined by Fisher’s exact test and Chi-square test. For quantification by immunofluorescence staining, statistically significant difference was determined by Student’s t-test. For all statistical analyses, *p*-values < 0.05 (two-tailed) were deemed statistically significant. Bar graphs represent means +/-SEM.

## Acknowledgement

We are grateful to members of Dr. Aviv Regev’s laboratory (Broad Institute), especially Danielle Dionne, Lan Nguyen, Julia Waldman, Michael Cuoco, Michal Slyper, Christopher Rodman, Orr Ashenberg, Marcin Tabaka, and Orit Rozenblatt-Rosen, as well as Dr. Asaf Rotem (Dana-Farber Cancer Institute) and Dr. Michael Steinbaugh (Harvard Chan Bioinformatics Core), for assistance with scRNA-seq processing and analysis. We thank Dr. Roderick Bronson at Harvard Medical School for histopathology consultation, the Harvard Nikon Imaging Center for technical support in immunofluorescence imaging, and all members of the Brugge laboratory, especially Jason Zoeller, Jennifer Rosenbluth, Mackenzie Boedicker, and Nomeda Girnius, for their assistance in this study. This work was supported in part by funding from a Susan G. Komen Postdoctoral Fellowship (C.M.L.), a Croucher Postdoctoral Fellowship (C.M.L.), a NIH F31 Fellowship (G.K.G.), the Hope Funds Postdoctoral Fellowship (Y.O.), the Marsha Rivkin Center for Ovarian Cancer Research (Y.O.), the Chan Zuckerberg Initiative Donor-Advised Fund (grant number 2018-182734 to L.P.), an advised fund of Silicon Valley Community Foundation (L.P.), a National Human Genome Research Institute (NHGRI) Career Development Award (R00HG008399 to L.P.), the Susan G. Komen Breast Cancer Foundation (J.S.B.), and the Breast Cancer Research Foundation (J.S.B.).

## Figure Legends

**Supplementary figure 1.**
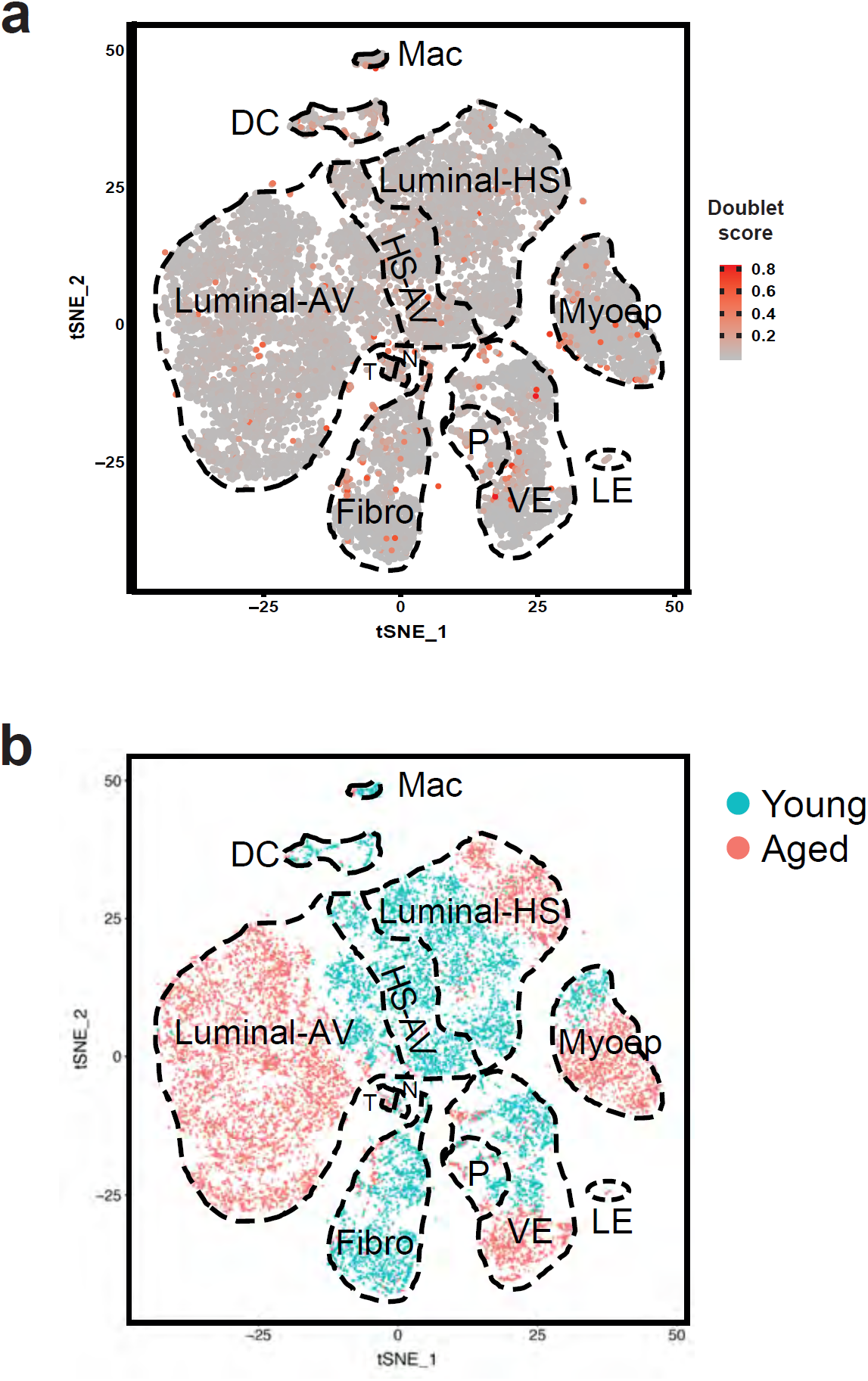
t-SNE plots related to Figure 1. **a** Predicted doublet scores in scRNA-seq data using Scrublet. A higher score indicates a greater likelihood of a cell event being a doublet. **b** t-SNE plot colored by age groups.

**Supplementary figure 2.**
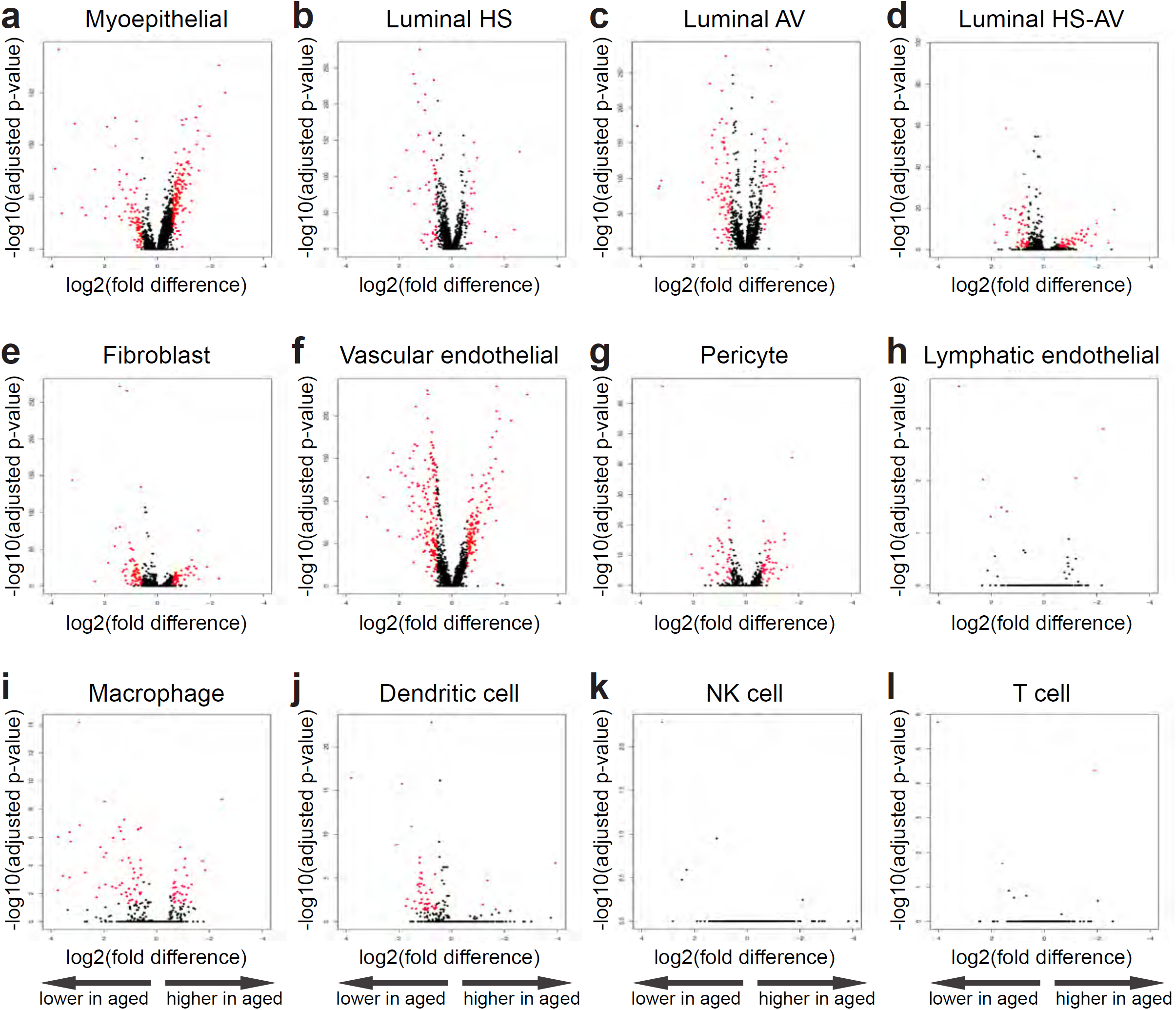
Overview of gene expression patterns in young and aged mammary glands. Volcano plots show expression levels of all detected genes in aged mammary glands (n = 4) compared to young mammary glands (n = 3) within the specified cell types. Genes with fold-difference above 1.5 and with adjusted *p*-value < 0.05 are highlighted in red.

**Supplementary figure 3.**
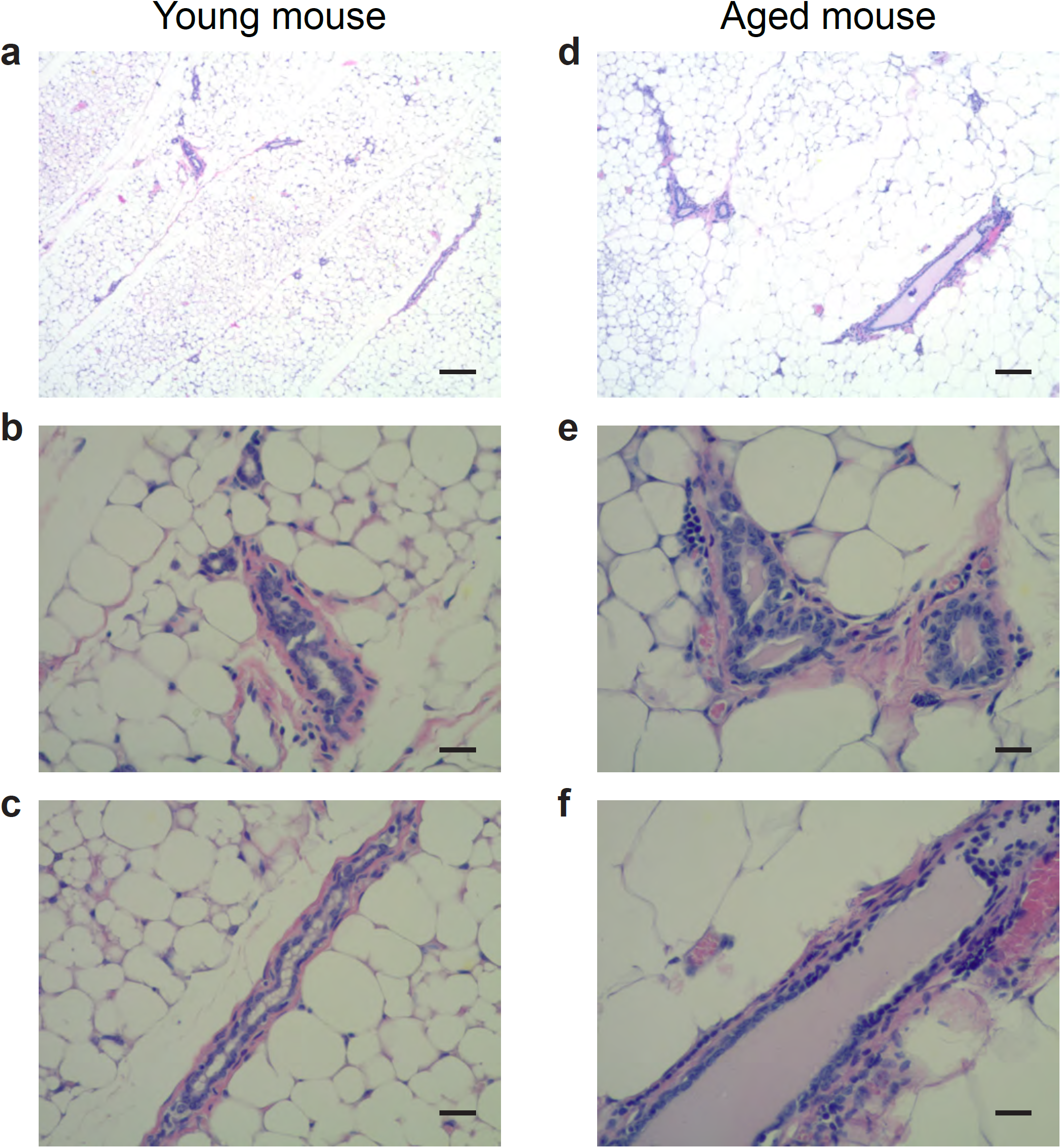
Representative H&E images of young (**a**-**c**) and aged (**d**-**f**) murine mammary glands at 4x (**a** and **d**) and 20x magnifications (**b, c, e**, and **f)**. Aged mammary glands show increased secretory material in alveoli (**e**) and dilated ducts (**f**). In **a** and **d**, scale bar = 100 *µ*m. In **b, c, e**, and **f**, scale bar = 20 *µ*m.

**Supplementary figure 4.**
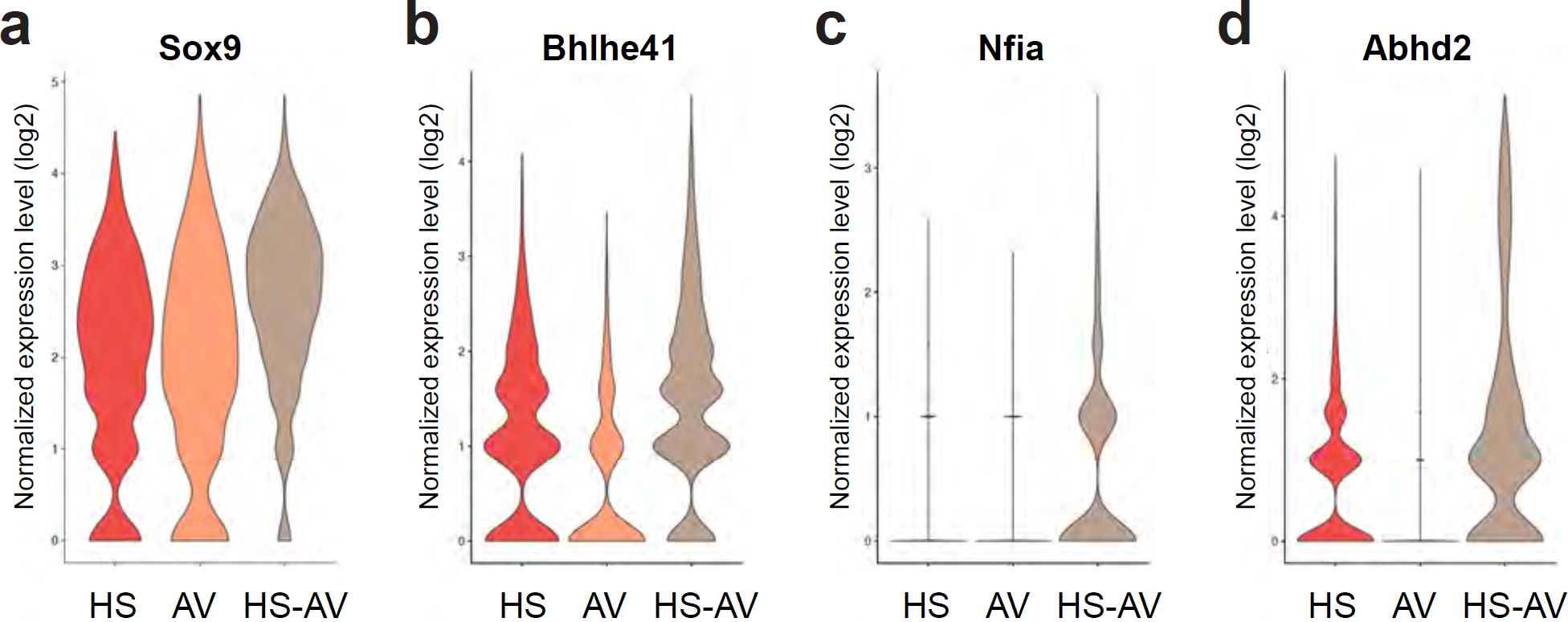
Select genes identified in Fig. 5a with expression enriched in HS-AV luminal cells relative to HS cells and AV cells.

**Supplementary figure 5.**
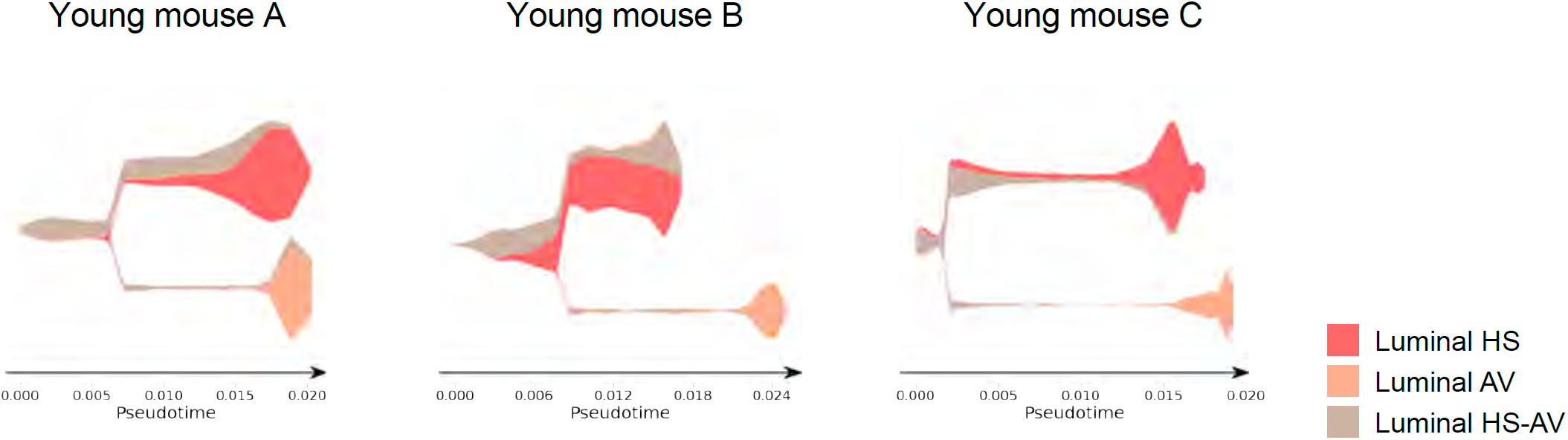
STREAM plots from lineage trajectory analysis of luminal cells in young mammary glands (n = 3 samples). Related to Fig. 5b.

**Supplementary figure 6.**
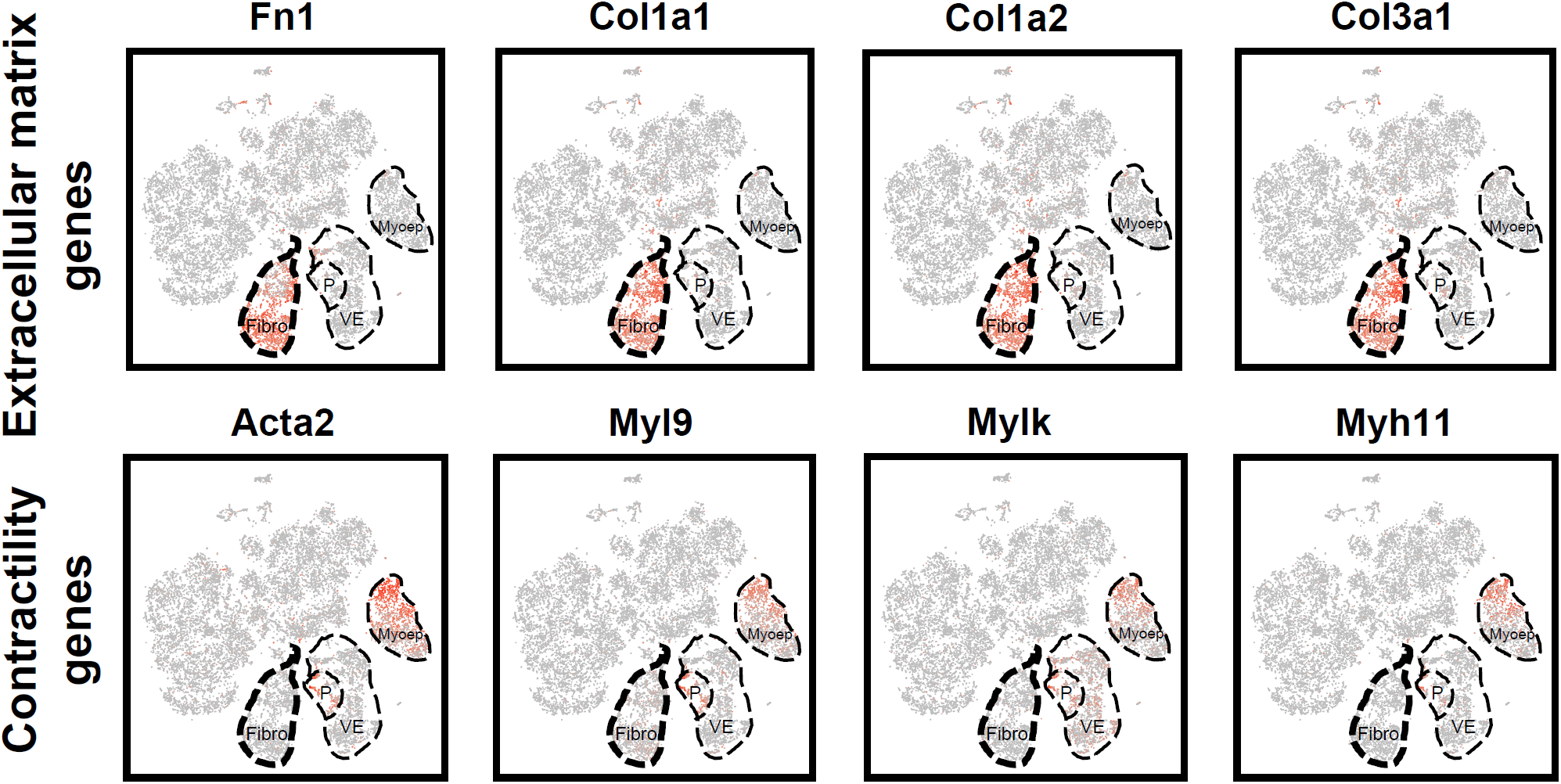
The fibroblasts captured in our scRNA-seq analysis of both young and aged mammary glands express characteristic extracellular matrix genes. However, they do not express contractility genes, in contrast to myoepithelial cells, pericytes, and vascular endothelial cells, which do express these markers. The lack of contractility gene expression in these fibroblasts indicates that they are unlikely to be myofibroblasts.

**Supplementary table 1.** Cell type-specific gene marker signatures. Related to Fig. 1g.

**Supplementary table 2.** Chi-square test results for all per-sample pairwise comparisons of relative proportions of cell types. Related to Fig. 2a. The higher Chi-square values for pairwise per-sample comparisons across age groups than within age groups reflect the age-dependent differences in cell type compositions.

**Supplementary table 3.** Percent abundance of cell types in young and aged mammary glands. Related to Fig. 2a-d.

**Supplementary table 4.** Differentially expressed genes in aged mammary glands compared to young mammary glands within each cell type. Related to Fig. 3c, 4e, 4f, 6i, 6l, and 7i.

**Supplementary table 5.** HS-AV cell markers. Related to Fig. 5a.

