## Supplementary table 2 for "Aging-associated alterations in the mammary gland revealed by single-cell RNA sequencing"

**Supplementary table 2. Chi-square test results for all per-sample pairwise comparisons of relative proportions of cell types**

| Chi-square | Young (A) | Young (B) | Young (C) | Aged (A) | Aged (B) | Aged (C) | Aged (D) |
|---|---|---|---|---|---|---|---|
| Young (A) | | | | | | | |
| Young (B) | 193 | | | | | | |
| Young (C) | 591 | 757 | | | | | |
| Aged (A) | 1745 | 1469 | 2435 | | | | |
| Aged (B) | 2604 | 2126 | 3525 | 162 | | | |
| Aged (C) | 2613 | 2031 | 3262 | 238 | 370 | | |
| Aged (D) | 1899 | 1577 | 2816 | 124 | 77 | 228 | |

All p-values < 0.0001; df = 8
