## Supplementary table 3 for "Aging-associated alterations in the mammary gland revealed by single-cell RNA sequencing"

### Supplementary table 3. Percent abundance of cell types in young and aged mammary glands

#### Relative abundance across all cell types

| | Young (A) | Young (B) | Young (C) | Aged (A) | Aged (B) | Aged (C) | Aged (D) |
|---|---|---|---|---|---|---|---|
| Myoepithelial | 7.9% | 3.2% | 1.5% | 12.3% | 17.9% | 6.0% | 16.6% |
| Luminal-HS | 19.9% | 25.9% | 30.8% | 11.8% | 16.3% | 6.6% | 11.3% |
| Luminal-AV | 10.3% | 7.3% | 11.4% | 57.3% | 53.6% | 75.0% | 54.8% |
| Luminal-HS-AV | 17.6% | 17.5% | 17.7% | 0.3% | 0.5% | 0.1% | 0.3% |
| Fibroblast | 19.2% | 12.1% | 27.6% | 5.6% | 1.1% | 0.7% | 0.9% |
| Vascular endothelial | 13.7% | 24.9% | 3.8% | 9.4% | 8.0% | 8.9% | 13.9% |
| Pericyte | 4.1% | 4.5% | 1.1% | 1.2% | 1.3% | 1.7% | 1.4% |
| Lymphatic endothelial | 0.3% | 0.1% | 0.2% | 0.1% | 0.1% | 0.4% | 0.3% |
| Immune | 7.0% | 4.5% | 5.8% | 2.0% | 1.0% | 0.6% | 0.5% |
| TOTAL | 100% | 100% | 100% | 100% | 100% | 100% | 100% |

#### Relative abundance across epithelial cell types

| | Young (A) | Young (B) | Young (C) | Aged (A) | Aged (B) | Aged (C) | Aged (D) |
|---|---|---|---|---|---|---|---|
| Myoepithelial | 14.2% | 5.9% | 2.5% | 15.0% | 20.3% | 6.8% | 20.0% |
| Luminal-HS | 35.7% | 48.0% | 50.1% | 14.5% | 18.4% | 7.5% | 13.6% |
| Luminal-AV | 18.4% | 13.6% | 18.6% | 70.1% | 60.7% | 85.6% | 66.0% |
| Luminal-HS-AV | 31.6% | 32.5% | 28.8% | 0.4% | 0.6% | 0.1% | 0.4% |
| TOTAL | 100% | 100% | 100% | 100% | 100% | 100% | 100% |

#### Relative abundance across stromal cell types

| | Young (A) | Young (B) | Young (C) | Aged (A) | Aged (B) | Aged (C) | Aged (D) |
|---|---|---|---|---|---|---|---|
| Fibroblast | 43.3% | 26.2% | 71.7% | 30.5% | 9.8% | 5.9% | 5.1% |
| Vascular endothelial | 31.0% | 54.0% | 10.0% | 51.5% | 69.0% | 72.5% | 82.0% |
| Pericyte | 9.4% | 9.8% | 2.9% | 6.5% | 11.4% | 14.1% | 8.0% |
| Lymphatic endothelial | 0.7% | 0.3% | 0.5% | 0.8% | 1.1% | 3.0% | 1.7% |
| Immune | 15.7% | 9.8% | 15.0% | 10.8% | 8.7% | 4.5% | 3.1% |
| TOTAL | 100% | 100% | 100% | 100% | 100% | 100% | 100% |

#### Relative abundance across immune cell types

| | Young (A) | Young (B) | Young (C) | Aged (A) | Aged (B) | Aged (C) | Aged (D) |
|---|---|---|---|---|---|---|---|
| Macrophage | 40.8% | 18.7% | 8.2% | 20.9% | 15.8% | 41.7% | 27.3% |
| Dendritic cell | 51.8% | 66.7% | 79.2% | 37.2% | 15.8% | 8.3% | 9.1% |
| Natural killer cell | 5.7% | 4.0% | 9.1% | 11.6% | 7.9% | 8.3% | 9.1% |
| T cell | 1.8% | 10.7% | 3.5% | 30.2% | 60.5% | 41.7% | 54.5% |
| TOTAL | 100% | 100% | 100% | 100% | 100% | 100% | 100% |
